# Lipid Dynamics and Organization Around Voltage-Gated Sodium Channels: A Coarse-Grained Perspective

**DOI:** 10.1101/2025.08.04.668480

**Authors:** Abhinav Srivastava, Paweł Chodnicki, Jacek Czub, Vincenzo Carnevale

## Abstract

Lipid–ion channel interactions play a critical role in channel function and membrane structural organization. Despite this importance, the mechanisms behind the rearrangement of lipids around ion channels are still unclear. To investigate this, we conducted coarse grained (CG) molecular dynamics simulations of voltage gated sodium ion channels (NavAB) embedded in a ternary lipid bilayer composed of 1,2-dilinoleoyl-sn-glycero-3-phosphocholine (DIPC), 1,2-dipalmitoyl-sn-glycero- 3-phosphocholine (DPPC), and cholesterol (CHOL) at varying CHOL concentrations (6.62%, 17.62% and 30.00%). By analyzing lipid organization and membrane structure, we examined how membrane composition and channel state (activated and resting) influence lipid redistribution near the channel interface. Our key finding is a pronounced preference for DIPC for the channel vicinity, observed consistently for all CHOL concentrations and channel states. Our simulations reveal that hydrophobic mismatch dictates lipid sorting near NavABs. The hydrphobic thickness of the channel favors flexible DIPC lipids, which are packed efficiently around it, while excluding thicker DPPC lipids. This exclusion drives DPPC and cholesterol to form ordered domains farther from the channel interface. Mixing entropy analysis supports local lipid de-mixing near the channel, aligning with the emergence of phase-separated domains. Notably, the hydrophobic thickness of NavAB remained stable and in close agreement with the experimental values, indicating that lipid-specific properties drive reorganization near the channel. Overall, our findings demonstrate that hydrophobic mismatch is a key driver of lipid reorganization and domain formation around ion channels, regardless of CHOL concentration or channel conformational state.

## Introduction

Ion channels are typically multimeric proteins embedded in the plasma membrane. They contain a pore that spans the membrane and undergo conformational changes to regulate its opening and closing. These pores can open and close in response to various types of signals, such as the binding of specific ligands(1–3), mechanical stimuli such as membrane tension(4, 5) and changes in voltage across the membrane(6– 9). When open, ions traverse the channel in a single-file manner, moving either into or out of the cell. Ion channels mediate the selective movement of ions across the cell mem-branes, which is a fundamental mechanism that enables electrical excitability in muscle cells and governs most forms of electrical signaling in the nervous system. Voltage-gated ion channels have voltage sensor domains that detect changes in membrane potential. These channels initiate and propagate action potentials in response to membrane depolarization, playing a crucial role in the functioning of muscle, nerve, and other electrically excitable cells(10–16). Voltage-gated sodium channels (NavABs) initiate in response to membrane depolarization and preferentially conduct Na^+^ over other monovalent or divalent cations. The activated state refers to a conformation of the ion channel in which the pore is open, allowing the ions to pass through, while the pore is closed in the resting state, thereby preventing the ion conduction.

Lipids can interact directly with ion channels through lipidprotein interactions, modulating the conformational changes that drive the opening and closing of ion channels during the gating process(17–20). Mostly, lipid-ion channel interaction occurs when ion channel gating charges interact with lipid head groups, thereby making stable electrostatic interactions with the negatively charged phosphate moiety of lipids(21, 22). The dependence of lipids on voltage-gated sodium channels was reported by investigating the activity of three bacterial sodium channel homologues (NaChBac, NavMs, and NavSp) with varying lipid compositions. Each channel was found to show a unique lipid preference, with NavMs and NavSp favoring different negatively charged lipids, while NaChBac showed minimal variation. Structural and molecular modeling studies revealed that lipid specificity arises mainly from interactions with channel voltage sensing subdomains(23). For example, potassium ion channels in their open, intermediate, and closed states were embedded in five distinct lipid bilayers and investigated using both experimental methods and molecular dynamics simulations. The study revealed that changes in the lipid composition can alter the electrostatic environment surrounding the ion channels, affecting the TM potential. This, in turn, can indirectly modulate the channel activity and function(24). In addition to the well-established role of lipids in membrane structure, recent studies have highlighted the influence of lipid phase state on ion channel function. The interaction between lipid rafts and voltage-gated ion channels can lead to clustering, cooperativity and long-term memory effects, potentially mod-ulating channel gating and signaling(25). Lipids modulate ions through four mechanisms: acting as coupling elements, facilitating migrating mechanisms, serving as fixing points, or functioning as molecular wedges. This lipid modulation contributes to overall regulatory mechanisms, presenting a potential avenue to identify new pharmacological targets for drug design by disrupting lipid-channel interactions(26).

Previous studies have extensively investigated lipid-channel interactions and their role in modulating ion channel function. However, the effects of CHOL concentration on the organization of lipids and the formation of domains around NavABs in both the activated and resting states remain unclear. To address this gap, we performed CG simulations of NavABs embedded in ternary lipid mixtures with different CHOL concentrations. We systematically characterized lipid organization and domain formation near NavABs, revealing that DIPC preferentially localized at the channel interface because of hydrophobic compatibility, while DPPC forms ordered domains away from the channel. The lipid organization is governed by the intrinsic phase behavior of the lipid mixture and occurs independently of the CHOL concentration or conformational state of the channel. The consistent hydrophobic thickness of NavABs throughout the simulations confirmed protein structural stability. Together, these findings deepen our understanding of how lipid-driven forces, rather than channel activity, shape membrane organization.

### Simulation methodology

All CG simulations were performed using GROMACS- 2023.3 patched with PLUMED 2(27–34). We used Martini 2.2 force fields for NavAB, lipids and CHOL(35–37).

Electrostatic interactions were calculated using the reaction field method with a cutoff of 1.1 nm, consistent with Martini parameters(38). van der Waals interactions were considered between all beads within a 1.1 nm range, using a cutoff potential. The Verlet neighbor search algorithm(39) was employed to enhance computational efficiency. The temperature was maintained at 303.15 K using the velocity rescale thermostat(40) with a coupling constant of 1 ps and pressure was maintained at 1 bar using semi-isotropic Parrinello-Rahman pressure coupling(41) with a time constant of 12 ps and compressibility of 3.0*X*10^*−*4^ bar^-1^. All simulations ran with a time step of 20 fs for 5 *µ*s per system. The systems consisted of NavAB channels in activated and resting states embedded in ternary lipid bilayers consisting DIPC, DPPC and CHOL. The cryo-EM structures of the activated and resting state NavABs were obtained from the OPM and RCSB PDB database respectively (PDB IDs: 6P6Y and 6P6W)(42). We prepared our systems using the CHARMM-GUI Martini Maker(43–45). The bilayer composition included equimolar proportions of DIPC and DPPC with CHOL added at low (6.62%), medium (17.62%) and high (30.00%) molar concentrations. We selected DIPC as a representative unsaturated lipid commonly used in CG simulations, associated with liquid disordered (L_d_) that promotes membrane thinning and fluidity due to its unsaturated acyl chains(46). In contrast, DPPC represents a typical saturated lipid that forms a liquid-ordered (L_O_) phase that contributes to membrane thickening and tighter lipid packing(47, 48). CHOL modulates the structural properties of both the L_d_ and L_O_ domains and influences the lipid-protein interactions by altering the order of the membrane. CG representations of lipids and CHOL are shown in Figure1. The functional groups of DIPC and DPPC include NC3, PO4, GL1 and GL2 corresponding to the moeities of choline, phosphate and glycerol. Acyl chains are mapped to C1A–C4A and C1B–C4B beads. For CHOL, ROH corresponds to the hydroxyl group, R1–R5 to the sterol rings, and C1, C2 as the tails. We began our simulations by executing all eight steps of energy minimization and equilibration as specified by the CHARMM-GUI Builder. During equilibration, NavAB channels were restrained with a force constant of 1000 kJ mol^-1^ nm^-2^ and lipid restraints were progressively reduced from 200 to 10 kJ mol^-1^. For multichannel systems, we replicated the systems in the X and Y directions. An additional energy minimization step was subsequently performed to eliminate any potential atomic overlaps. To avoid aggregation between NavABs, we implemented flat-bottom harmonic walls using PLUMED(33, 34). The lateral position (X,Y) of each channel’s center of mass (COM) was used as a positional reference. A 50 kJ mol^-1^ nm^-2^ restraint was applied allowing individual channels to move freely within defined regions, thus avoiding clustering (see FigureS1 of the supplementary material).

**Fig. 1.**
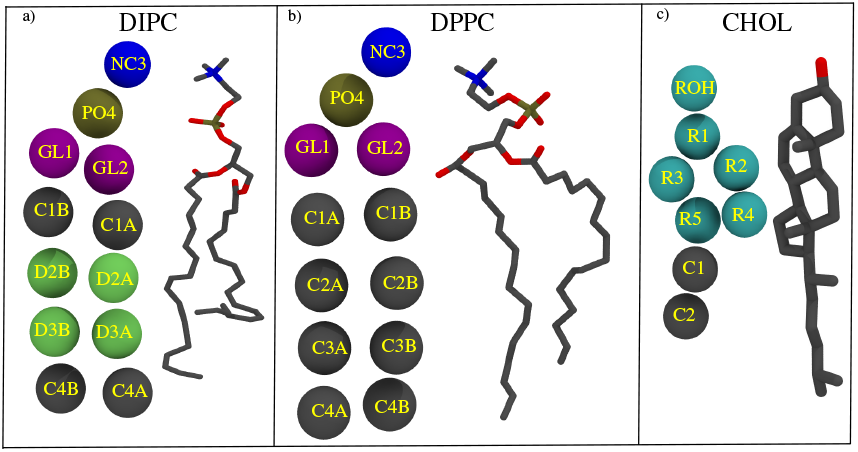
CG and all-atom representations of a) DIPC, b) DPPC, and c) CHOL, illustrating CG bead mappings to functional groups.

To investigate whether lipid reorganization arises from interchannel interactions or is an intrinsic response to the presence of a channel, we also simulated a single-channel system with the same lipid composition and CHOL concentration. In this setup, the channel was restrained in the center of the membrane using the same spatial restraints as in the multichannel system (see FigureS2 of the supplementary material). This allowed a direct comparison of the lipid properties in single-channel versus multiple-channel environments. Trajectory analysis and visualization were performed using the python packages MDAnalaysis(49, 50) and FREUD(51, 52).

## Results

We first analyze lipid distributions around NavAB channels in membranes with varying CHOL concentrations. Figure2 shows representative snapshots of the channel in low,medium, and high CHOL mixtures, highlighting the distinct lipid arrangements in the activated and resting states. These snapshots clearly indicate a consistent pattern of phase separation between the DIPC and DPPC lipids. DIPC shows a tendency to localize around NavABs thereby forming visible clusters at the channel-lipid interface. This trend persists for all CHOL concentrations and in both conformational states of the channel, suggesting that a local membrane environment near the channel favors the presence of DIPC. However, DPPCs are largely excluded from these regions and populate the surrounding membrane, often forming larger continuous domains. This apparent segregation likely reflects the underlying physiochemical differences between lipids such as bilayer thickness, fluidity or packing.

**Fig. 2.**
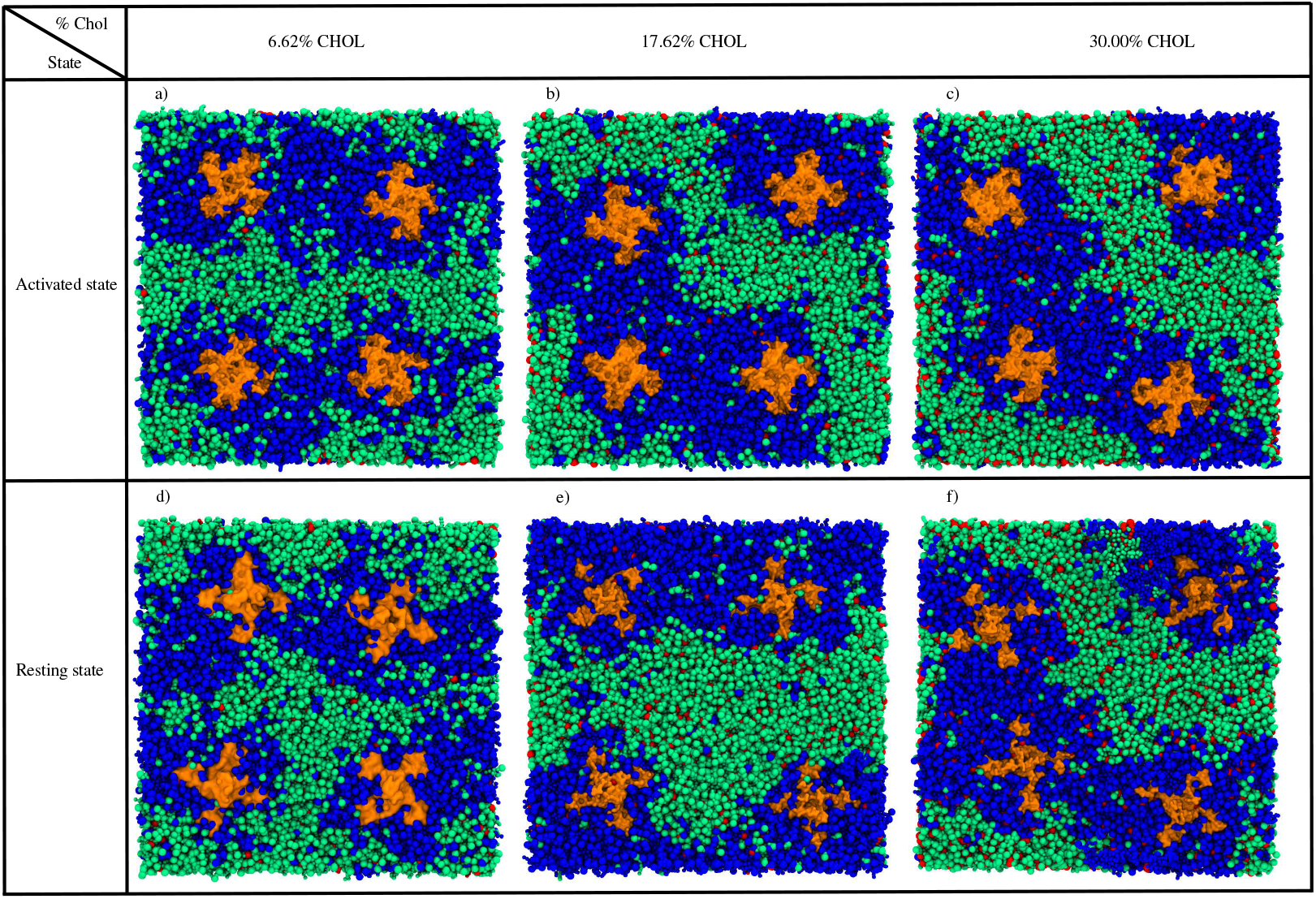
Snapshots of NavABs in a ternary mixture of DIPC/DPPC/CHOL at three different CHOL concentrations in both activated and resting states. Panels a), b) and c) show NavAB in the activated state at low, medium and high CHOL concentrations respectively while panels d), e) and f) represent NavAB in the resting state. All the snapshots are captured at 5*µ*s. Color Code: Blue (DIPC), Green (DPPC), Red (CHOL), and Orange (NavABs surface). Lipid CG bonds are visualized using a TCL script from the Martini webpage in VMD(53–55).

### Radial distribution of lipids around NavABs

To quantify the spatial organization of lipids around the NavABs, we computed the radial distribution function (g(r)) of DIPC and DPPC with respect to each NavABs (Figure3). The COM of each channel was used as reference and the PO4 bead of DIPC and DPPC was selected to reflect the importance of headgroup interactions in lipid-protein interactions(56, 57). This analysis captures how lipid density varies as a function of distance from NavAB, enabling identification of lipid enrichment or depletion zones. The top and bottom panels of Figure 3 show g(r) profiles for the activated and resting states respectively. The profiles were averaged over all four channels. In the activated state at low CHOL,the DIPC exhibits a strong localization around NavABs. This is evident from a pronounced peak in g(r) at ≈1.5 nm suggesting direct interactions between DIPC and NavABs. The emergence of other peaks indicates a significant lipid enrichment near NavABs. Additional small undulations in the g(r) at longer distances suggest the presence of long-range interactions between DIPC and NavABs. In contrast, the g(r) of DPPC is nearly flat, indicating a lack of preferential association implying that DIPC is excluded from the channel environment, likely residing in more ordered membrane domains farther away from NavABs. At medium CHOL, DIPC remains largely unchanged, with a persistent and well-defined first peak around 1.5 nm, a height similar to that observed at low CHOL. This suggests that even with an increase in CHOL concentration, DIPC continues to maintain strong interactions with NavAB in the activated state. For DPPC, g(r) remains low and relatively featureless, although a minor upward slope is observed at larger distances. At high CHOL, DIPC continues to show a strong affinity for NavABs. Interestingly, in the activated state at high CHOL concentrations, the primary g(r) peak for DIPC is lower compared to other CHOL concentrations. While this difference falls within the range of the error bars shown in the plot, the trend is consistent for other systems. This may suggest that an increase in the concentration of CHOL indicates subtle changes in membrane organization that marginally reduce the accessibility of DIPC to the channel surface, although strong DIPC-channel association persists overall.

**Fig. 3.**
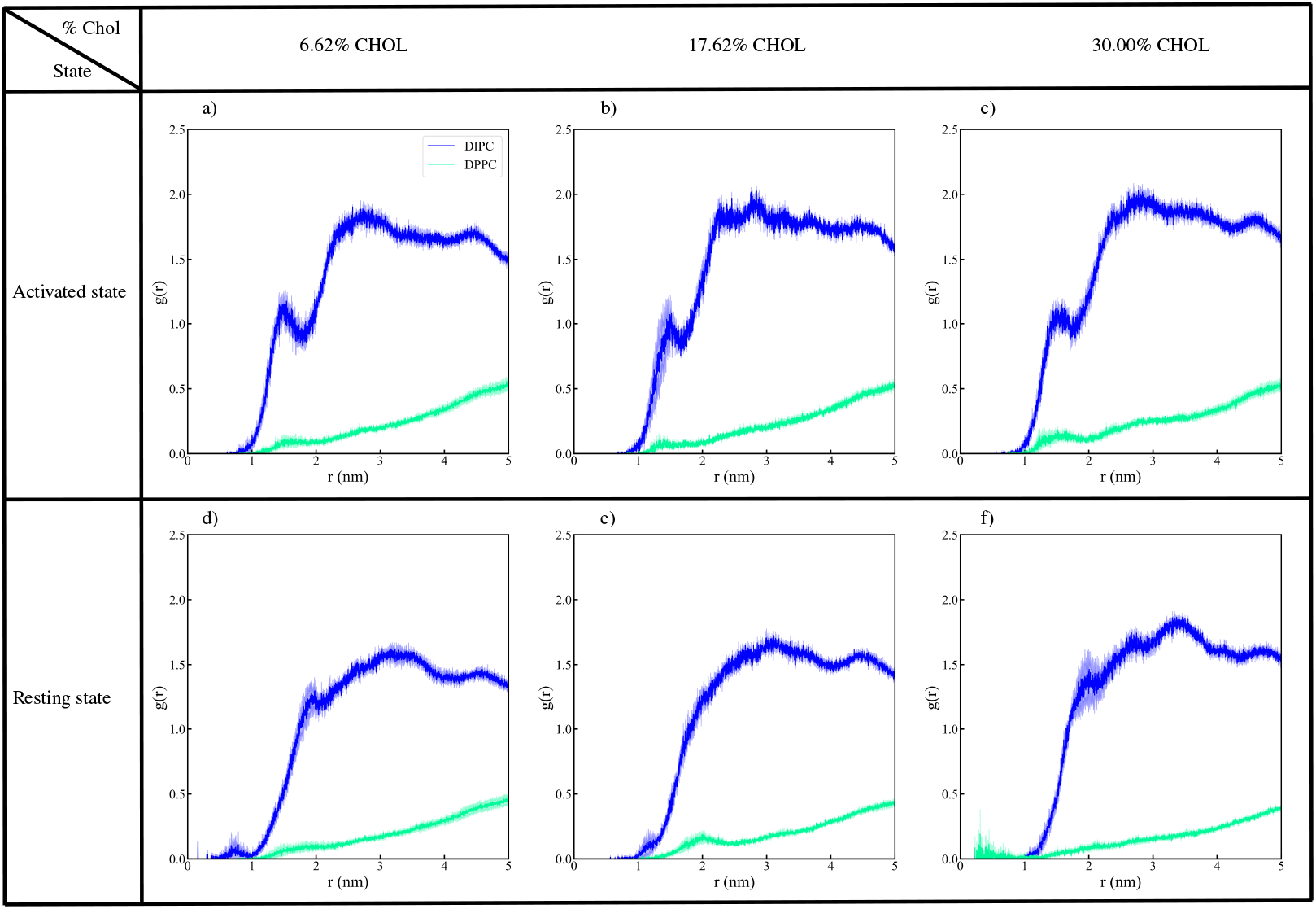
Radial distribution function, *g*(*r*), of DIPC and DPPC relative to NavAB channels at low, medium and high CHOL concentrations. Solid blue and green lines represent DIPC and DPPC respectively, with shaded regions indicating error bars. The *g*(*r*) were computed individually for each of the four channels and averaged to provide a comprehensive representation of lipid distribution around NavABs

In the resting state, DIPC remains enriched near NavAB with g(r) peaks centered around ≈1.8 nm. Compared with those in the activated state, these peaks are slightly broader and shifted, which may reflect changes in the lipid packing or local membrane organization. For all CHOL concentrations, DPPC exhibits flat g(r) profiles with near-zero values close to the channel, thereby reaffirming its exclusion from the immediate lipid environment. These trends may arise from conformational differences in NavAB that modulate lipid accessibility with DIPC consistently forming the dominant annular shell. At medium CHOL, the DIPC’s g(r) profile lacks sharp peaks and shows a broader elevation, indicating a less pronounced interaction with NavAB in comparison to the low CHOL concentration, though some enrichment remains. The absence of distinct radial structuring may suggest reduced ordering of DIPC around the channel. DPPC continues to show a mostly flat profile (a shallow peak near ≈2.0 nm, consistent with redistribution effects likely driven by CHOL-induced domain formation rather than specific channel association. This gradual increase lacks a defined peak, indicating that there is no specific enrichment near NavAB. Instead, the weak DPPC profile can be interpreted as a redistribution effect, which likely reflects the formation of the CHOL-induced domains.

To assess whether differences in lipid composition are associated with changes in the structural dynamics of NavAB, we calculated the root mean square deviation (RMSD) of NavABs in both the activated and resting states for all CHOL concentrations. RMSD provides a quantitative measure of structural fluctuation over time, offering insight into how stable or flexible the channel remains under different membrane conditions. The RMSD shown in FigureS3 of the supplementary material reveals that the activated state consistently exhibits lower RMSD values than the resting state for all CHOL concentrations. Both states plateaued within 1 *µ*s indicating equilibration on the simulation timescale. Interestingly, with an increase in the CHOL concentration, we observe an increase in the RMSD in both conformations, suggesting that higher CHOL concentrations may induce additional conformational adjustments. When considered alongside the g(r) profiles, the RMSD results reveal that structural fluctuations are consistently lower in the activated state compared to the resting state for all CHOL concentrations. RMSDs increase slightly with an increase in CHOL in both conformations, which may reflect changes in membrane-channel interactions. These trends highlight how the lipid composition and the bilayer environment modulate the conformational dynamics of NavAB channels over the simulation timescales.

### Analysis of lipid clustering patterns

To dissect the spatial heterogeneity of lipid distributions around NavABs, clustering was employed to quantify lipid domain formation, providing a more detailed structural assessment of DIPC and DPPC localization patterns at low, medium and high CHOL concentrations in activated and resting states. Although g(r) captures average lipid preferences near the channel, they do not account for lateral organization or domain-level segregation across the membrane surface. Therefore, we applied clustering to directly visualize and quantify lateral lipid domains and assess how these are modulated by CHOL concentration and channel states. We chose the upper leaflet of the membrane for our clustering analysis. To assign lipids to individual leaflets, we first calculate the bilayer midplane using the COM of all PO4 beads along the z-axis. Lipids whose PO4 beads z-coordinates were above the midplane were classified as belonging to the upper leaflet, and those below as part of the lower leaflet. Subsequently, we performed clustering using the density-based spatial clustering of applications with noise (DBSCAN) algorithm(58), where a predefined neighborhood distance (2.5 nm and a minimum number of lipids (15) were used to determine the clusters. Lipids that do not meet the clustering threshold were identified as noise. Larger sample sizes typically lead to the formation of fewer, but larger clusters due to stricter density requirements, while smaller sample sizes may result in more fragmented clusters or noise because the algorithm is able to detect smaller clustering with fewer lipids.

Figure4 shows the spatial organization of DIPC and DPPC lipids taken from the simulation frame where the maximum number of DIPC clusters was observed, along with its corresponding DPPC distribution. The figure illustrates results for low, medium and high CHOL concentrations in both activated and resting states. The clustering pattern reflects the preferential localization of DIPC near NavABs. Around NavABs, DIPC molecules form large clusters, contributing to well-separated domains, whereas in regions farther from the channel, they form smaller, more scattered clusters, particularly at lower CHOL concentrations, leading to an overall higher number of observed clusters. In contrast, DPPC, which avoids channel-proximal regions, forms fewer, larger clusters in the remaining membrane space, suggesting a spatial reorganization in which DIPC dominates the regions near NavABs, while DPPC form larger domains farther from the channel. The influence of CHOL is evident both in the cluster counts and in the spatial organization. CHOL is known to enhance membrane rigidity by increasing bilayer thickness(59– 61). Furthermore, CHOL preferentially interacts with DPPC due to its higher affinity for saturated lipids(62, 63), promoting the formation of more ordered and condensed DPPC-rich domains. As CHOL integrates within these domains, it stabilizes larger DPPC aggregates, contributing to the observed reduction in cluster numbers for both lipids as the CHOL concentration increases. The effect remains consistent in both the activated and resting states, reflecting the role of CHOL in promoting lipid condensation and stabilizing domain formation.

**Fig. 4.**
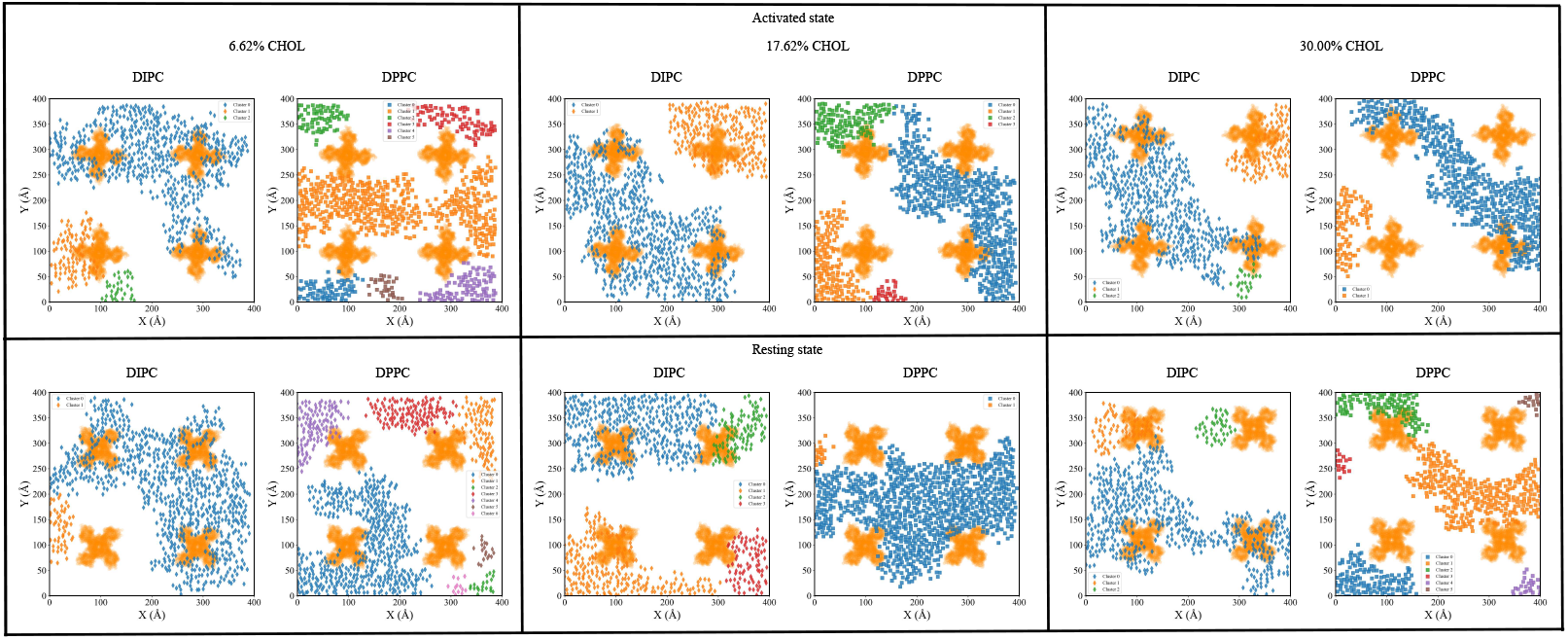
Spatial arrangement of DIPC and DPPC clusters at low, medium and high CHOL concentrations in both activated and resting states. Top row (activated state): The spatial snapshots depict the largest DIPC cluster (colored based on number or cluster) and the corresponding spatially equivalent DPPC region for comparison at each CHOL concentration. Bottom row (resting state): Similar snapshots are shown for the restig state, illustrating the spatial distribution of the largest DIPC cluster and its DPPC counterpart. The clusters are computed at a sample size of 15.

In the activated state, DIPC clusters closely associate with the channel regions, while DPPC forms larger, more peripheral clusters away from NavABs. In the resting state, DIPC clusters remain near the channel but appear to be more dispersed compared to the activated state because of NavAB’s increased flexibility. This increased flexibility, as evidenced by RMSD, prevents DIPC from forming tightly packed clusters around NavAB, unlike in the activated state, where the channel’s greater structural stability of the channel supports the more compact DIPC clustering near NavABs. The higher structural stability in the activated state may create a more favorable hydrophobic environment for DIPC, reinforcing its localization near the channel and promoting tighter clustering. Meanwhile, the sustained flexibility of the resting state disrupts this organization, leading to more scattered, less cohesive DIPC clusters despite the lipid’s continued preference for channel-proximal regions.

### Voronoi tessellation

Voronoi tessellation is crucial for studying the lipid organization around NavABs, as it partitions the membrane into regions that define the spatial territory of each lipid molecule. This geometry-based approach provides information on lipid distribution and local crowding effects. In this analysis, each lipid headgroup serves as a generating point, and the resulting Voronoi diagram delineates regions corresponding to individual lipid molecules, offering a detailed view of local packing and spatial organization. The method enables us to examine how lipid environments vary, with distinct spatial patterns emerging in the presence of NavAB channels.

For tessellation, we selected DIPC and DPPC lipids in the upper leaflet and used the Freud package to compute the Voronoi, considering PO4 bead of each lipid as a Voronoi seed. The algorithm divides the bilayer plane into regions, assigning each point to its closest lipid head group, effectively describing the spatial territory of each lipid. To complement this, a hexbin plot was generated with a grid size of 60, partitioning the bilayer into hexagonal bins and counting the number of DIPC molecules within each hexagon. To ensure that low-density regions are still represented, we included bins containing at least one DIPC molecule, ensuring a comprehensive view of both densely packed and sparsely populated areas.

Figure5 shows the Voronoi tessellation that illustrates the spatial distribution of DIPC (blue) and DPPC (green) lipids around NavABs (orange) at low, medium and high CHOL concentrations. At low CHOL, DIPC forms a relatively uniform distribution throughout the membrane but exhibits a clear tendency to cluster near NavABs. This pattern suggests that DIPC’s unsaturated, thinner hydrophobic tails create a favorable interaction with the channel surface, likely driven by a hydrophobic mismatch. In contrast, DPPC appears to be randomly distributed, with no strong preference for channel regions. The resting state exhibits a similar distribution pattern, though the DIPC density is slightly reduced near NavABs.

**Fig. 5.**
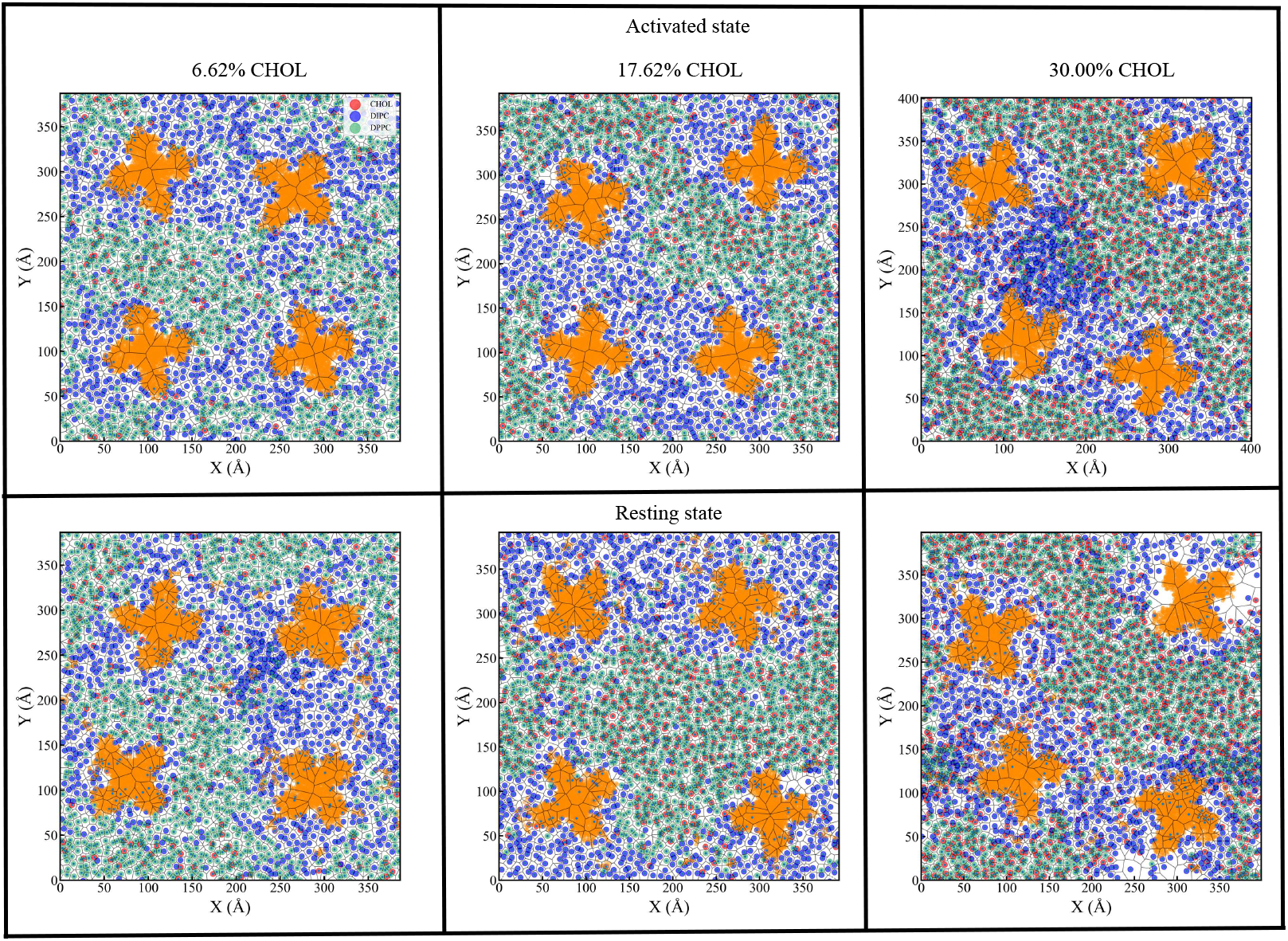
Voronoi tessellation plots of DIPC (blue) and DPPC (green) lipid densities around NavABs (orange) for low, medium and high CHOL concentrations. The top and bottom row show Voronoi tessellation of activated and resting state respectively. Color intensity indicates lipid density (scale bars on the right). These snapshots were taken at 5*µ*s.

In medium CHOL, a shift in the lipid organization becomes apparent. DIPC maintains its preference for the channel vicinity, but DPPC, supported by CHOL’s stabilizing effect on ordered phase, start to form larger patches in the non-channel regions. This emerging phase separation indicates a competitive balance between DIPC’s affinity for the channel and DPPC’s CHOL driven stabilization. In the resting state, this effect becomes more pronounced: DPPC forms an increasingly contiguous phase, thereby driving DIPC to smaller channel-adjacent domains. This behavior supports the idea that CHOL promotes the ordering of DPPC(64–68), gradually excluding DIPC from membrane regions.

At high CHOL, phase separation becomes more distinct. DPPC forms extensive ordered regions that dominate the membrane surface, while DIPC is progressively confined to small NavAB-proximal regions. In the activated state, DIPC forms dense clusters near the channel, reinforcing the role of hydrophobic mismatch in the sustainability of this interaction. The resting state exhibits an even starker separation, with the DPPC completely occupying the bulk membrane, and the DIPC almost entirely restricted to the immediate vicinity of NavABs. This observation aligns with the notion that CHOL preferentially associates with saturated lipids such as DPPC, amplifying domain formation and minimizing the presence of DIPC in non-NavAB regions.

### Hydrophobic mismatch

Our analysis of lipid localization and clustering revealed that DIPC preferentially localizes near NavABs, compared to DPPC. One possible physical mechanism underlying this preferential localization is a hydrophobic mismatch that can drive lipid reorganization by inducing membrane deformation or channel adaptation. This mismatch arises when the hydrophobic thickness of the membrane differs from that of the TM region of the channel. This discrepancy leads to energetic penalties that can be relieved by tilting or stretching the TM helices, local membrane thinning or thickening, or selective lipid partition near the channel interface(69). The hydrophobic mismatch was quantified by comparing the hydrophobic thickness of NavABs (defined as the average distance between hydrophobic residues at the upper and lower TM boundaries) with that of pure DIPC/CHOL and DPPC/CHOL bilayers for low, medium and high CHOL concentrations. To isolate intrinsic bilayer thickness and avoid local distortions caused by protein insertion, these membrane-only systems were simulated separately. As shown in Tables1 and 2, the calculated hydrophobic mismatch was consistently greater for the DPPC-rich membrane regions than for the DIPC-rich ones for all CHOL concentrations and in the activated and resting states. This suggests that DIPC’s preferential association with NavABs may be driven, at least in part, by its better hydrophobic matching, minimizing the energetic cost of lipid-channel interactions. In table1, we observe that for DIPC lipids, membrane thickness increased with increasing CHOL concentration, ranging from 3.61 nm (low CHOL) to 3.78 nm (high CHOL). Meanwhile, the TM thickness of NavAB remained relatively constant in the activated state (2.91 −2.88 nm) resulting in a gradually increasing hydrophobic mismatch from 0.70 to 0.90 nm. For DPPC, in L_o_ phase, the membranes were substantially thicker (4.24 *−*4.46 nm), leading to a more pronounced mismatch with NavAB (1.33 *−*1.57 nm).

**Table 1.** T_Protein_, T_membrane_, and hydrophobic mismatch (H.M) for DIPC/CHOL and DPPC/CHOL bilayers in the activated state for low, medium and high CHOL concentrations. All units are expressed in nm.

| Lipid | CHOL conc. | T<sub>Protein</sub> | T<sub>Membrane</sub> | H.M |
| --- | --- | --- | --- | --- |
| DIPC | Low | 2.91 | 3.61 | 0.70 |
|  | Medium | 2.92 | 3.69 | 0.77 |
|  | High | 2.88 | 3.78 | 0.90 |
| DPPC | Low | 2.91 | 4.24 | 1.33 |
|  | Medium | 2.92 | 4.36 | 1.44 |
|  | High | 2.88 | 4.46 | 1.57 |

**Table 2.** T_Protein_, T_membrane_ and H.M for DIPC/CHOL and DPPC/CHOL bilayers in the resting state for low, medium and high CHOL concentrations. All units are expressed in nm.

| Lipid | CHOL conc. | T<sub>Protein</sub> | T<sub>Membrane</sub> | H.M |
| --- | --- | --- | --- | --- |
| DIPC | Low | 2.73 | 3.61 | 0.88 |
|  | Medium | 2.78 | 3.69 | 0.90 |
|  | High | 2.72 | 3.78 | 1.06 |
| DPPC | Low | 2.73 | 4.24 | 1.51 |
|  | Medium | 2.78 | 4.36 | 1.58 |
|  | High | 2.72 | 4.46 | 1.75 |

In the resting state (Table2), the TM thickness of NavAB decreased slightly to 2.72 *−* 2.78 nm, further increasing the hy-drophobic mismatch in both lipid systems. For DIPC, mismatch values increased to 0.88 *−*1.06 nm while for DPPC, the mismatch became more severe, reaching 1.51 *−*1.75 nm. These data demonstrate that the hydrophobic mismatch increases consistently with CHOL concentrations and is consistently higher for DPPC than for DIPC for both activated and resting state NavABs.

Importantly, this increasing mismatch correlates with the observed structural stability of the channel, especially in the resting conformation. For example, in the resting state, where the mismatch is more pronounced, the channel exhibits a higher RMSD over time as compared to the activated state. This suggests that a larger mismatch imposes mechanical strain on the TM helices, potentially forcing the channel to adapt to a more flexible conformation in order to minimize the energetic cost associated with the mismatch. The observed lipid organization near the channel further supports this interpretation. DIPC, in L_d_ phase, preferentially localizes near the channel, while DPPC, L_o_ phase, is depleted at the channel interface. This mismatch likely contributes to both lipid sorting and the conformational dynamics of the channel(70).

### Entropy of mixing

The mixing entropy (S_Mix_) is an important thermodynamic parameter that quantifies the disorder in a multi-component system, increasing as the system transitions from a fully demixed state to a mixed state(71, 72). To quantify S_Mix_ of lipids in activated and resting state, we apply the approach proposed by Straub et al.(73). S_Mix_ is given by the following equation:

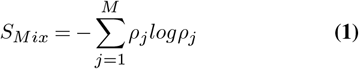

where *ρ*_*j*_ represents the mole fraction of each component *j* in a system with *M* components(74). The mole fraction *ρ*_*j*_ is defined as 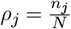, where n_j_ is the number of molecules of the component *j* (DIPC, DPPC and CHOL in our simulations) and *N* is total molecules. Thus, *ρ*_*j*_ captures the proportion of each component in the overall mixture. For a well-mixed blend in which the mole fraction of the components are similar, S_Mix_ will be relatively high. In contrast, when the system is phase separated or more segregated, S_Mix_ decreases as one or more components become dominant. To quantify the compositional heterogeneity of our system, we calculate the S_Mix_ of the upper leaflets for low, medium and high concentrations of CHOL in both activated and resting states (Figure6). We choose the upper leaflet because the bilayer leaflets can evolve asymmetrically due to protein interactions, cholesterol distribution, and local membrane stresses(75). Calculating S_Mix_ for the entire bilayer averages these differences, potentially masking the biologically relevant leaflet-specific behavior. Therefore, focusing on the upper leaflet preserves spatial resolution and better captures changes in lipid composition that can influence protein function.

In the activated state at low CHOL, S_Mix_ starts at ≈1.00 (ideal mixed system) and gradually decreases to ≈0.80 suggesting limited demixing and the retention of moderate mixing during the course of simulations (Figure6). In contrast, systems in medium CHOL show a more rapid decrease in entropy, with S_Mix_ falling below 0.70 at the end of simulation. This indicates a more substantial segregation of lipid components, likely driven by an increase in the CHOL concentration, thereby altering the local lipid environment. At high CHOL, S_Mix_ drops to values below 0.65 indicating that there is a strong compositional demixing, consistent with the formation of well-defined domains in which specific lipid types become locally enriched while others are excluded. In particular, in this region, the entropy reaches a near plateau within the first 2 *−*3 *µ*s suggesting that large-scale rearrangements and domain stabilization occur relatively early and persist throughout the remainder of the simulation.

**Fig. 6.**
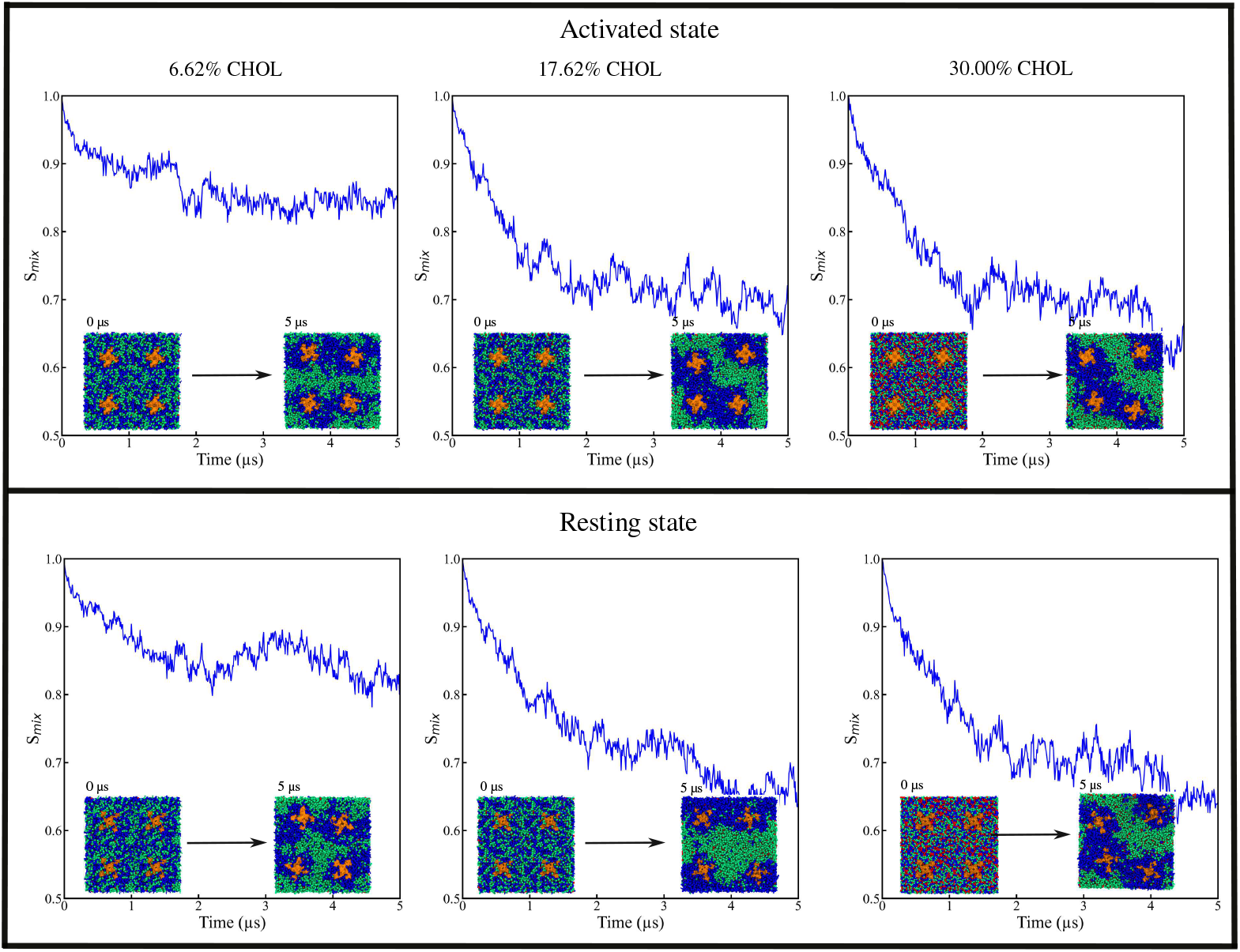
Time evolution of S_mix_ for DIPC, DPPC and CHOL in activated (top row) and resting state NavABs (bottom row) at low, medium and high CHOL concentrations. Each plot include system snapshots at initial (0*µ*s) and final simulation time (5*µ*s for NavABs.

When comparing activated and resting conformational states, S_Mix_ trends are similar for each CHOL concentration, while minor differences in S_Mix_ can be observed at certain time instants, there is no systematic or significant deviation between the two states. This indicates that the conformational state of NavABs does not play a major role in modulating lipid demixing at the global level. Instead, CHOL concentration plays an important role in influencing the degree of lipid mixing and segregation.

Our results reveal that increasing CHOL concentration leads to a consistent and concentration-dependent reduction in mixing entropy, pointing to enhanced lateral segregation of lipid species in the membrane. This suggests that the membrane transitions from a relatively mixed state at low CHOL to a more compositionally heterogeneous organization at higher CHOL levels, regardless of the structural state of the embedded protein.

## Conclusions

In this study, we present a detailed investigation of lipid-channel interactions in heterogeneous bilayers, focusing on how hydrophobic mismatch drives lipid sorting and domain formation around the sodium ion channels. Using a combina-tion of complementary analyses including radial distribution functions, clustering, Voronoi tessellation, hydrophobic mismatch and entropy of mixing, we reveal how lipid composition and protein conformational state influence membrane organization.

Radial distribution function analysis shows a consistent and preferential accumulation of DIPC lipids near the channel surface, while DPPC is largely excluded from the immediate vicinity of NavAB. This pattern persists for both activated and resting states and is robust for low, medium and high CHOL concentrations. These observations reflect the thinner, more disordered nature of DIPC, which provides better hydrophobic matching with the channel’s TM region. In contrast, DPPC’s thicker and more ordered profile leads to poor hydrophobic compatibility, promoting its segregation into CHOL-enriched domains farther from the protein. Building on this, the root mean square deviation analysis shows that NavAB adopts a more stable conformation in the activated state as compared to the resting state, particularly at high CHOL concentrations. This likely shows hydrophobic compatibility between the DIPC and the activated-state conformation, leading to strong lipid-channel interactions and reduced structural fluctuations. In the resting state, the mismatch between DDPC-rich regions and the channel surface may result in less favorable packing, which contributes to greater conformational flexibility. These state-dependent differences underscore how the lipid and channel state can mutually influence membrane-channel dynamics.

Voronoi tessellation and clustering analyses shed light on the lateral lipid organization. DIPC forms persistent interfacial clusters around the channel, while DPPC and CHOL segregate into more ordered domains. Voronoi-based spatial partitioning highlights how lipid packing geometries adjust to accommodate the channel footprint, further reinforcing the role of local lipid rearrangement in minimizing hydrophobic mismatch.

The entropy of mixing analysis shows a consistent decrease in S_Mix_ over time for low, medium and high CHOL concentrations and both channel states, indicating progressive lipid demixing and domain formation. In particular, DIPC becomes enriched near the channel, whereas DPPC and CHOL segregate into more ordered regions farther away. These observations highlight the role of hydrophobic mismatch and CHOL-mediated lipid preferences in driving the lateral membrane reorganization around NavAB.

In general, our findings establish hydrophobic mismatch as a central determinant of lipid sorting and membrane heterogeneity around the embedded ion channel. The channel does not activatedly dictate membrane organization; rather, it participates in shaping the surrounding lipid landscape by virtue of its hydrophobic profile. Lipids reorganize in response, minimizing energetic penalties and forming distinct spatial domains. This lipid-driven behavior has important implications not just for channel function, but also for understanding how membrane composition and channel structure co-evolve. Our results open new directions for studying channel-lipid interactions in more complex environments, such as asym-metric bilayers, different ion channels. As lipidomics and biology continue to converge, integrating physical insights will be crucial for decoding the role of membrane in cellular signaling and channel function.

## Author Contributions

V.C designed the research. A.S and P.C. carried the simulations, analyzed the data and wrote in-house python scripts to analyze the trajectories. V.C and J.C helped in analysis and writing the manuscript.

## Acknowledgments

The authors express their gratitude to the Gdańsk University of Technology for their support under the ‘Excellence Initiative Research University (IUDB)’ program. The computations were carried out using High Performance Cluster facilities of the Center of Informatics at the Tricity Academic Supercomputer and Network (Poland) and Poland’s High Performance Infrastructure PLGrid (ACK Cyfronet Ares). The authors would also like to thank Mateusz Kogut for his valuable help.

## Notes

### Competing Interest Statement

The authors have declared no competing interest.

### Funding

- Nobelium Joining Gdansk Tech Research Community: DEC-43/2023/IDUB/I.1

## Bibliography

1. Ferdinand Hucho and Christoph Weise. Ligand-gated ion channels. Angewandte Chemie International Edition, 40:3100–3116, 2001.

2. Joseph W. Lynch and Peter H. Barry. Ligand-Gated Ion Channels: Permeation and Activation1, pages 335–367. Springer New York, 2007.

3. Yanggan Wang, Deeptankar DeMazumder, and Joseph A. Hill. Chapter 7 - ionic fluxes and genesis of the cardiac action potential. In Muscle, pages 67–85. Academic Press, 2012.

4. Bradley Akitake, Andriy Anishkin, and Sergei Sukharev. The “dashpot” mechanism of stretch-dependent gating in mscs. The Journal of general physiology, 125:143–154, 2005.

5. Charles D Cox, Toshifumi Nomura, CS Ziegler, Anthony Keith Campbell, Kenneth Taylor Wann, and Boris Martinac. Selectivity mechanism of the mechanosensitive channel mscs revealed by probing channel subconducting states. Nature communications, 4:2137, 2013.

6. Anatoly Dryga, Suman Chakrabarty, Spyridon Vicatos, and Arieh Warshel. Realistic simulation of the activation of voltage-gated ion channels. Proceedings of the National Academy of Sciences, 109:3335–3340, 2012.

7. Gildas Loussouarn and Mounir Tarek. Editorial: Molecular mechanisms of voltage-gating in ion channels. Frontiers in Pharmacology, 12, 2021.

8. Wenlei Ye, Hongtu Zhao, Yaxin Dai, Yingdi Wang, Yu hua Lo, Lily Yeh Jan, and Chia-Hsueh Lee. Activation and closed-state inactivation mechanisms of the human voltage-gated kv4 channel complexes. Molecular Cell, 82:2427–2442.e4, 2022.

9. Huiwen Chen, Zhanyi Xia, Jie Dong, Bo Huang, Jiangtao Zhang, Feng Zhou, Rui Yan, Yiqiang Shi, Jianke Gong, Juquan Jiang, Zhuo Huang, and Daohua Jiang. Structural mechanism of voltage-gated sodium channel slow inactivation. Nature Communications, 15: 3691, 2024.

10. Bert Hille. Ion Channels of Excitable Membranes. Sinauer Associates, Sunderland, MA, 3rd edition, 2001.

11. A. L. Hodgkin and A. F. Huxley. Resting and action potentials in single nerve fibres. The Journal of Physiology, 104:176–195, 1945.

12. A. L. Hodgkin and A. F. Huxley. A quantitative description of membrane current and its application to conduction and excitation in nerve. The Journal of Physiology, 117:500–544, 1952.

13. William A. Catterall. Structure and function of voltage-gated ion channels. Annual Review of Biochemistry, 64:493–531, 1995.

14. William A. Catterall. Ion channel voltage sensors: Structure, function, and pathophysiology. Neuron, 67:915–928, 2010.

15. William A Catterall. From ionic currents to molecular mechanisms: The structure and function of voltage-gated sodium channels. Neuron, 26:13–25, 2000.

16. Christopher A. Ahern, Jian Payandeh, Frank Bosmans, and Baron Chanda. The hitchhiker’s guide to the voltage-gated sodium channel galaxy. Journal of General Physiology, 147:1– 24, 2015.

17. Avia Rosenhouse-Dantsker, Dolly Mehta, and Irena Levitan. Regulation of Ion Channels by Membrane Lipids, pages 31–68. John Wiley Sons, Ltd, 2012. ISBN 9780470650714.

18. Julio F Cordero-Morales and Valeria Vásquez. How lipids contribute to ion channel function, a fat perspective on direct and indirect interactions. Current Opinion in Structural Biology, 51:92–98, 2018.

19. Anna L. Duncan, Wanling Song, and Mark S.P. Sansom. Lipid-dependent regulation of ion channels and g protein–coupled receptors: Insights from structures and simulations. Annual Review of Pharmacology and Toxicology, 60:31–50, 2020.

20. Andrea Saponaro and Marco Lolicato. Editorial: The key role of lipids in the regulation of ion channels. Frontiers in Physiology, 13, 2022.

21. Luis G Cuello, D Marien Cortes, and Eduardo Perozo. Molecular architecture of the kvap voltage-dependent k+ channel in a lipid bilayer. Science, 306:491–495, 2004.

22. Anthony Lee. Lipid interactions with ion channels. Future Lipidology, 1:103–114, 2006.

23. Nazzareno D’Avanzo, Emily C. McCusker, Andrew M. Powl, Andrew J. Miles, Colin G. Nichols, and B. A. Wallace. Differential lipid dependence of the function of bacterial sodium channels. PLOS ONE, 8:1–10, 2013.

24. Marina A. Kasimova, Mounir Tarek, Alexey K. Shaytan, Konstantin V. Shaitan, and Lucie Delemotte. Voltage-gated ion channel modulation by lipids: Insights from molecular dynamics simulations. Biochimica et Biophysica Acta (BBA) - Biomembranes, 1838:1322–1331, 2014.

25. Antonio Suma, Daniel Sigg, Seamus Gallagher, Giuseppe Gonnella, and Vincenzo Carnevale. Ion channels in critical membranes: Clustering, cooperativity, and memory effects. PRX Life, 2:013007, 2024.

26. José A. Poveda, A. Marcela Giudici, M. Lourdes Renart, Andrés Morales, and José M. González-Ros. Towards understanding the molecular basis of ion channel modulation by lipids: Mechanistic models and current paradigms. Biochimica et Biophysica Acta (BBA) - Biomembranes, 1859:1507–1516, 2017.

27. EJ Dijkstra S Achterop H Bekker, HJC Berendsen, D Van der Spoel R Vondrumen, A Sijbers, H Keegstra, and MKR Renardus. Gromacs - a parallel computer for molecular-dynamics simulations. In Physics Computing ‘92, pages 252–256. World Scientific Publishing, 1993.

28. H.J.C. Berendsen, D. van der Spoel, and R. van Drunen. Gromacs: A message-passing parallel molecular dynamics implementation. Computer Physics Communications, 91:43– 56, 1995.

29. E. Lindahl, B. Hess, and D. van der Spoel. Gromacs 3.0: a package for molecular simulation and trajectory analysis. Journal of Molecular Modeling, 7:306–317, 2001.

30. David Van Der Spoel, Erik Lindahl, Berk Hess, Gerrit Groenhof, Alan E. Mark, and Herman J. C. Berendsen. Gromacs: Fast, flexible, and free. Journal of Computational Chemistry, 26:1701–1718, 2005.

31. Berk Hess, Carsten Kutzner, David van der Spoel, and Erik Lindahl. Gromacs 4: Algorithms for highly efficient, load-balanced, and scalable molecular simulation. Journal of Chemical Theory and Computation, 4:435–447, 2008.

32. Sander Pronk, Szilárd Páll, Roland Schulz, Per Larsson, Pär Bjelkmar, Rossen Apostolov, Michael R. Shirts, Jeremy C. Smith, Peter M. Kasson, David van der Spoel, Berk Hess, and Erik Lindahl. Gromacs 4.5: a high-throughput and highly parallel open source molecular simulation toolkit. Bioinformatics, 29:845–854, 2013.

33. Massimiliano Bonomi, Davide Branduardi, Giovanni Bussi, Carlo Camilloni, Davide Provasi, Paolo Raiteri, Davide Donadio, Fabrizio Marinelli, Fabio Pietrucci, Ricardo A. Broglia, and Michele Parrinello. Plumed: A portable plugin for free-energy calculations with molecular dynamics. Computer Physics Communications, 180:1961–1972, 2009.

34. Gareth A. Tribello, Massimiliano Bonomi, Davide Branduardi, Carlo Camilloni, and Giovanni Bussi. Plumed 2: New feathers for an old bird. Computer Physics Communications, 185:604–613, 2014.

35. Luca Monticelli, Senthil K. Kandasamy, Xavier Periole, Ronald G. Larson, D. Peter Tieleman, and Siewert-Jan Marrink. The martini coarse-grained force field: Extension to proteins. Journal of Chemical Theory and Computation, 4:819–834, 2008.

36. Semen O. Yesylevskyy, Lars V. Schäfer, Durba Sengupta, and Siewert J. Marrink. Polarizable water model for the coarse-grained martini force field. PLOS Computational Biology, 6:1–17, 2010.

37. Djurre H. de Jong, Gurpreet Singh, W. F. Drew Bennett, Clement Arnarez, Tsjerk A. Wassenaar, Lars V. Schäfer, Xavier Periole, D. Peter Tieleman, and Siewert J. Marrink. Improved parameters for the martini coarse-grained protein force field. Journal of Chemical Theory and Computation, 9:687–697, 2013.

38. Ilario G. Tironi, René Sperb Paul E. Smith, and Wilfred F. van Gunsteren. A generalized reaction field method for molecular dynamics simulations. The Journal of Chemical Physics, 102:5451–5459, 1995.

39. Szilárd Páll and Berk Hess. A flexible algorithm for calculating pair interactions on simd architectures. Computer Physics Communications, 184:2641–2650, 2013.

40. Giovanni Bussi, Davide Donadio, and Michele Parrinello. Canonical sampling through velocity rescaling. The Journal of Chemical Physics, 126:014101, 2007.

41. M. Parrinello and A. Rahman. Polymorphic transitions in single crystals: A new molecular dynamics method. Journal of Applied Physics, 52:7182–7190, 1981.

42. Goragot Wisedchaisri, Lige Tonggu, Eedann McCord, Tamer M. Gamal El-Din, Liguo Wang, Ning Zheng, and William A. Catterall. Resting-state structure and gating mechanism of a voltage-gated sodium channel. Cell, 178(4):993–1003.e12, 2019.

43. Sunhwan Jo, Taehoon Kim, Vidyashankara G. Iyer, and Wonpil Im. Charmm-gui: A web-based graphical user interface for charmm. Journal of Computational Chemistry, 29(11):1859–1865, 2008.

44. Yifei Qi, Helgi I. Ingólfsson, Xi Cheng, Jumin Lee, Siewert J. Marrink, and Wonpil Im. Charmm-gui martini maker for coarse-grained simulations with the martini force field. Journal of Chemical Theory and Computation, 11:4486–4494, 2015.

45. Pin-Chia Hsu, Bart M. H. Bruininks, Damien Jefferies, Paulo Cesar Telles de Souza, Jumin Lee, Dhilon S. Patel, Siewert J. Marrink, Yifei Qi, Syma Khalid, and Wonpil Im. Charmm-gui martini maker for modeling and simulation of complex bacterial membranes with lipopolysaccharides. Journal of Computational Chemistry, 38:2354–2363, 2017.

46. Erwin London. How principles of domain formation in model membranes may explain ambiguities concerning lipid raft formation in cells. Biochimica et Biophysica Acta (BBA) - Molecular Cell Research, 1746:203–220, 2005.

47. J. Hjort Ipsen, G. Karlström, O.G. Mourtisen, H. Wennerström, and M.J. Zuckermann. Phase equilibria in the phosphatidylcholine-cholesterol system. Biochimica et Biophysica Acta (BBA) - Biomembranes, 905:162–172, 1987.

48. Todd PW McMullen, Ruthven NAH Lewis, and Ronald N McElhaney. Cholesterol– phospholipid interactions, the liquid-ordered phase and lipid rafts in model and biological membranes. Current opinion in colloid & interface science, 8:459–468, 2004.

49. Naveen Michaud-Agrawal, Elizabeth J. Denning, Thomas B. Woolf, and Oliver Beckstein. Mdanalysis: A toolkit for the analysis of molecular dynamics simulations. Journal of Computational Chemistry, 32(10):2319–2327, 2011.

50. R. J. Gowers, M. Linke, J. Barnoud, T. J. E. Reddy, M. N. Melo, S. L. Seyler, D. L. Dotson,J. Domanski, S. Buchoux, I. M. Kenney, and O. Beckstein. Mdanalysis: A python package for the rapid analysis of molecular dynamics simulations. In Proceedings of the 15th Python in Science Conference, pages 98–105. SciPy, 2016.

51. Bradley D. Dice, Vyas Ramasubramani, Eric S. Harper, Matthew P. Spellings, Joshua A. Anderson, and Sharon C. Glotzer. Analyzing particle systems for machine learning and data visualization with freud. In Proceedings of the 18th Python in Science Conference, pages 27–33, 2019.

52. Vyas Ramasubramani, Bradley D. Dice, Eric S. Harper, Matthew P. Spellings, Joshua A. Anderson, and Sharon C. Glotzer. Freud: A software suite for high throughput analysis of particle simulation data. Computer Physics Communications, 254:107275, 2020.

53. William Humphrey, Andrew Dalke, and Klaus Schulten. VMD – Visual Molecular Dynamics. Journal of Molecular Graphics, 14:33–38, 1996.

54. John Stone, Justin Gullingsrud, Paul Grayson, and Klaus Schulten. A system for interactive molecular dynamics simulation. In 2001 ACM Symposium on Interactive 3D Graphics, pages 191–194, 2001.

55. John Eargle, Dan Wright, and Zaida Luthey-Schulten. Multiple alignment of protein structures and sequences for vmd. Bioinformatics, 22:504–506, 2006.

56. Daniel Schmidt, Qiu-Xing Jiang, and Roderick MacKinnon. Phospholipids and the origin of cationic gating charges in voltage sensors. Nature, 444:775–779, 2006.

57. Yanping Xu, Yajamana Ramu, and Zhe Lu. Removal of phospho-head groups of membrane lipids immobilizes voltage sensors of k+ channels. Nature, 451:826–829, 2008.

58. Martin Ester, Hans-Peter Kriegel, Jorg Sander, and Xiaowei Xu. A density-based algorithm for discovering clusters in large spatial databases with noise. In kdd, volume 96, pages 226–231, 1996.

59. Norbert Kučerka, Jason D. Perlmutter, Jianjun Pan, Stephanie Tristram-Nagle, John Katsaras, and Jonathan N. Sachs. The effect of cholesterol on short- and long-chain monounsaturated lipid bilayers as determined by molecular dynamics simulations and x-ray scattering. Biophysical Journal, 95:2792–2805, 2008.

60. Frédérick de Meyer and Berend Smit. Effect of cholesterol on the structure of a phospholipid bilayer. Proceedings of the National Academy of Sciences, 106:3654–3658, 2009.

61. Tomasz Róg, Marta Pasenkiewicz-Gierula, Ilpo Vattulainen, and Mikko Karttunen. Ordering effects of cholesterol and its analogues. Biochimica et Biophysica Acta (BBA) - Biomembranes, 1788:97–121, 2009.

62. Xiaolian Xu and Erwin London. The effect of sterol structure on membrane lipid domains reveals how cholesterol can induce lipid domain formation. Biochemistry, 39:843–849, 2000.

63. Linda J. Pike. The challenge of lipid rafts. Journal of Lipid Research, 50:S323–S328, 2009.

64. J Hjort Ipsen, Ole G Mouritsen, and Martin J Zuckermann. Theory of thermal anomalies in the specific heat of lipid bilayers containing cholesterol. Biophysical journal, 56:661–667, 1989.

65. Oskar Engberg, Victor Hautala, Tomokazu Yasuda, Henrike Dehio, Michio Murata, J. Peter Slotte, and Thomas K.M. Nyholm. The affinity of cholesterol for different phospholipids affects lateral segregation in bilayers. Biophysical Journal, 111:546–556, 2016.

66. Matti Javanainen, Hector Martinez-Seara, and Ilpo Vattulainen. Nanoscale membrane domain formation driven by cholesterol. Scientific Reports, 7:1143, 2017.

67. Azadeh Alavizargar, Fabian Keller, Roland Wedlich-Söldner, and Andreas Heuer. Effect of cholesterol versus ergosterol on dppc bilayer properties: Insights from atomistic simulations. The Journal of Physical Chemistry B, 125:7679–7690, 2021.

68. Mohammadreza Aghaaminiha, Amir M. Farnoud, and Sumit Sharma. Quantitative relationship between cholesterol distribution and ordering of lipids in asymmetric lipid bilayers. Soft Matter, 17:2742–2752, 2021.

69. Armando J. de Jesus and Toby W. Allen. The determinants of hydrophobic mismatch response for transmembrane helices. Biochimica et Biophysica Acta (BBA) - Biomembranes, 1828:851–863, 2013.

70. Sarah L. Veatch and Sarah L. Keller. Separation of liquid phases in giant vesicles of ternary mixtures of phospholipids and cholesterol. Biophysical Journal, 85(5):3074–3083, 2003.

71. Peter Atkins and Julio De Paula. Atkins’ Physical Chemistry. Oxford University Press, Oxford, UK, 2006.

72. Marco Camesasca, Miron Kaufman, and Ica Manas-Zloczower. Quantifying fluid mixing with the shannon entropy. Macromolecular Theory and Simulations, 15:595–607, 2006.

73. George A. Pantelopulos, Tetsuro Nagai, Asanga Bandara, Afra Panahi, and John E. Straub. Critical size dependence of domain formation observed in coarse-grained simulations of bilayers composed of ternary lipid mixtures. The Journal of Chemical Physics, 147:095101, 2017.

74. Giovanni B. Brandani, Marieke Schor, Cait E. MacPhee, Helmut Grubmüller, Ulrich Zachariae, and Davide Marenduzzo. Quantifying disorder through conditional entropy: An application to fluid mixing. PLOS ONE, 8:1–8, 2013.

75. Georg Pabst and Sandro Keller. Exploring membrane asymmetry and its effects on membrane proteins. Trends in Biochemical Sciences, 49:333–345, 2024.

