## Supplementary information for "Lipid Dynamics and Organization Around Voltage-Gated Sodium Channels: A Coarse-Grained Perspective"

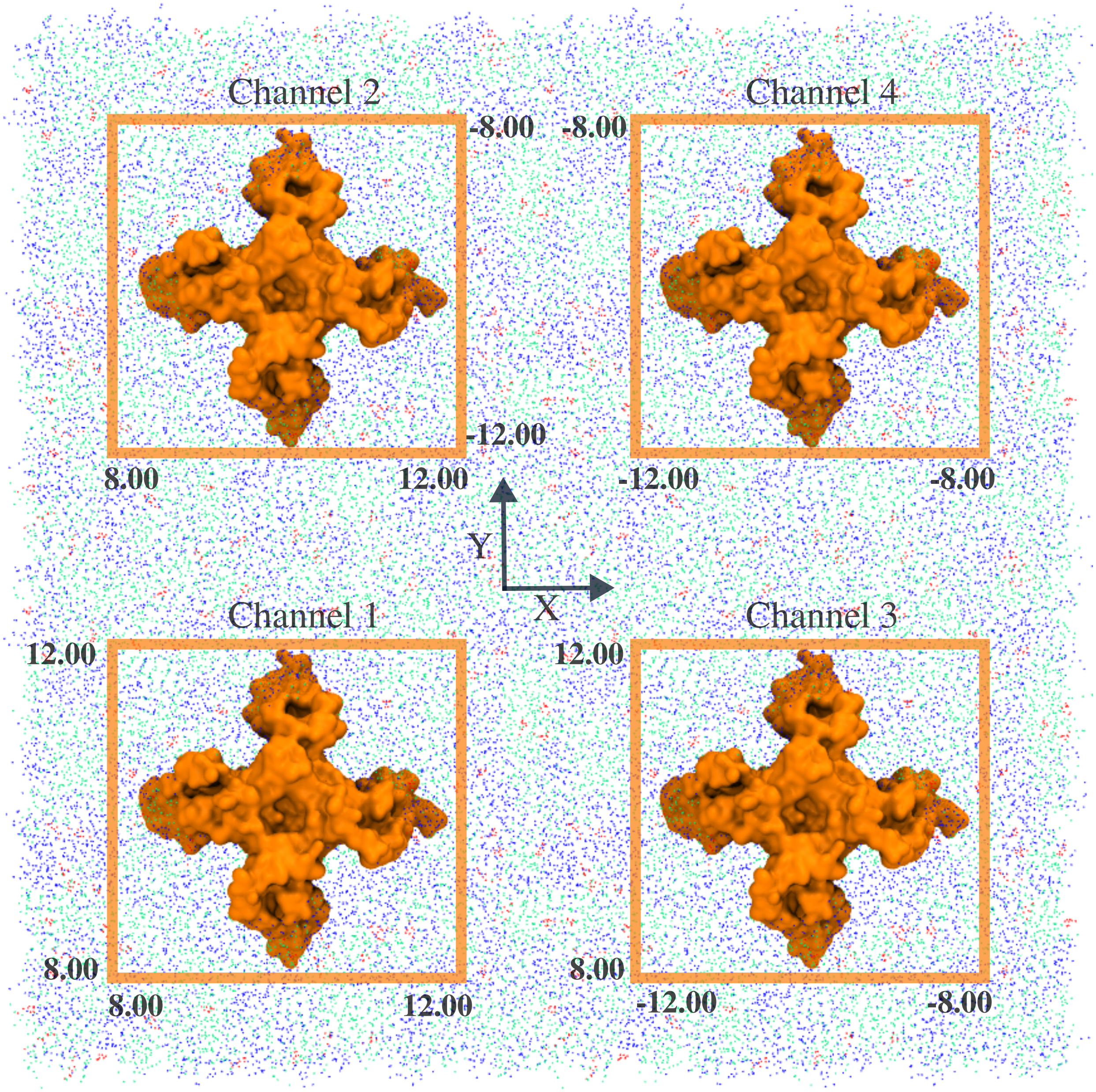


Figure S1: Representative snapshot of the simulation system showing the confinement regions for each NavAB. NavABs are numbered according to their respective confinement regions, with each channel restricted within specific ranges along X and Y directions. The confinement ensures that NavABs are isolated within defined regions. For clarity, lipids and CHOL are shown as transparent with the same color code as in Figure2 of the main text.

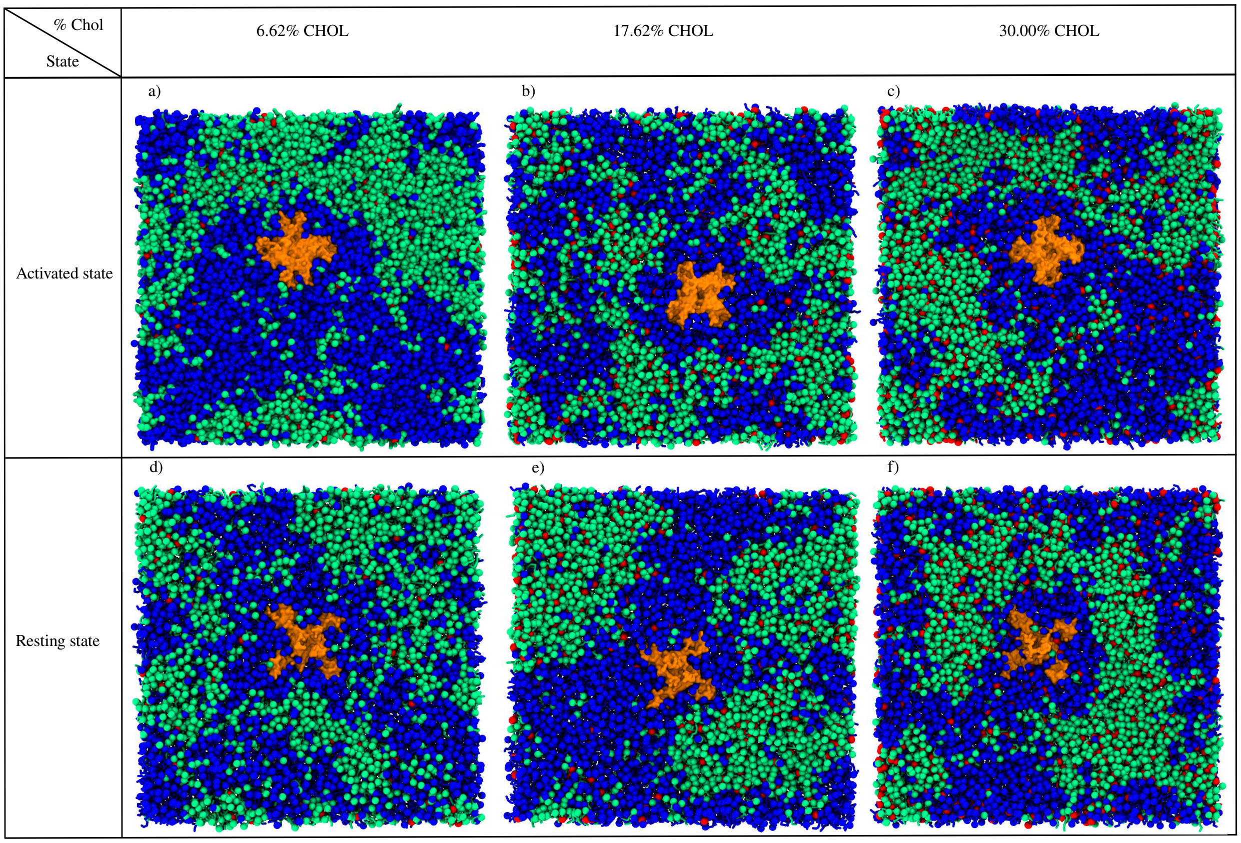


Figure S2: Snapshots of single NavAB channels in a ternary mixture of DIPC/DPPC/CHOL at low, medium and high CHOL concentrations. Panels a), b), and c) correspond to the activated state, while d), e), and f) show the resting state. All snapshots were taken at 5$\mu$s. Color code and visualization method are the same as in Figure2 of the main text.

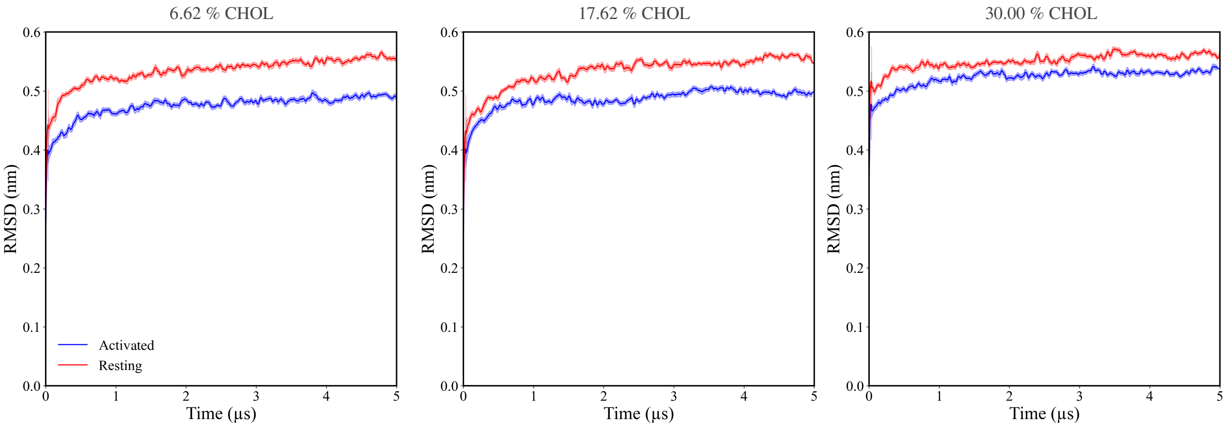


Figure S3: Root mean square deviation (RMSD) of NavAB channels in activated and resting states for low, medium, and high CHOL concentrations.

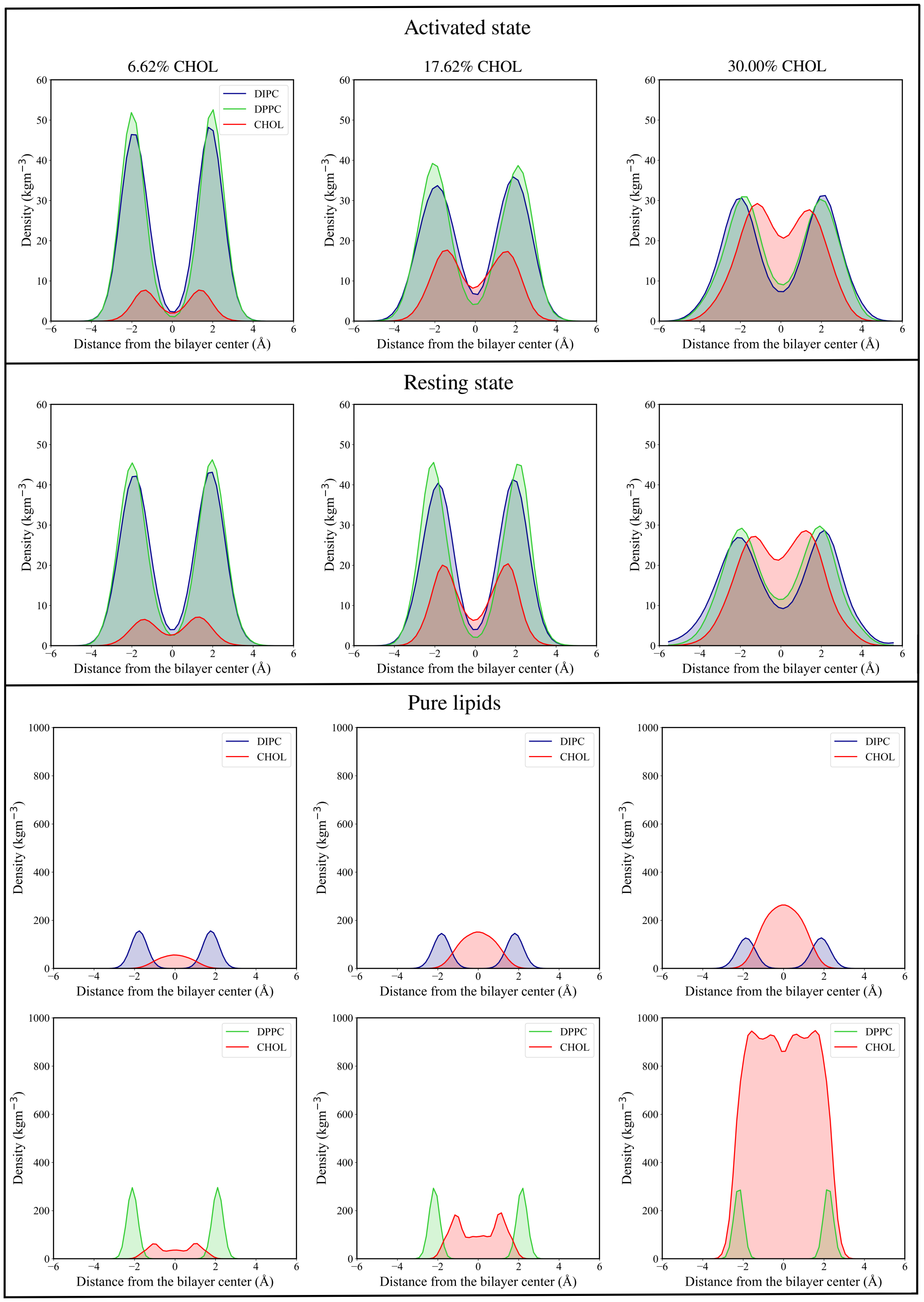


Figure S4: Density of DIPC, DPPC and CHOL at low, medium and high CHOL concentrations for activated and resting state NavABs. The first and second row showcase the densities in activated and resting states respectively. The third and forth row shows the densities of DIPC and DPPC pure lipids at low, medium and high CHOL concentrations respectively. In simulations, phosphate bead (PO4) for DIPC and DPPC, along with sterol bead (ROH) of CHOL, were taken to compute the densities.

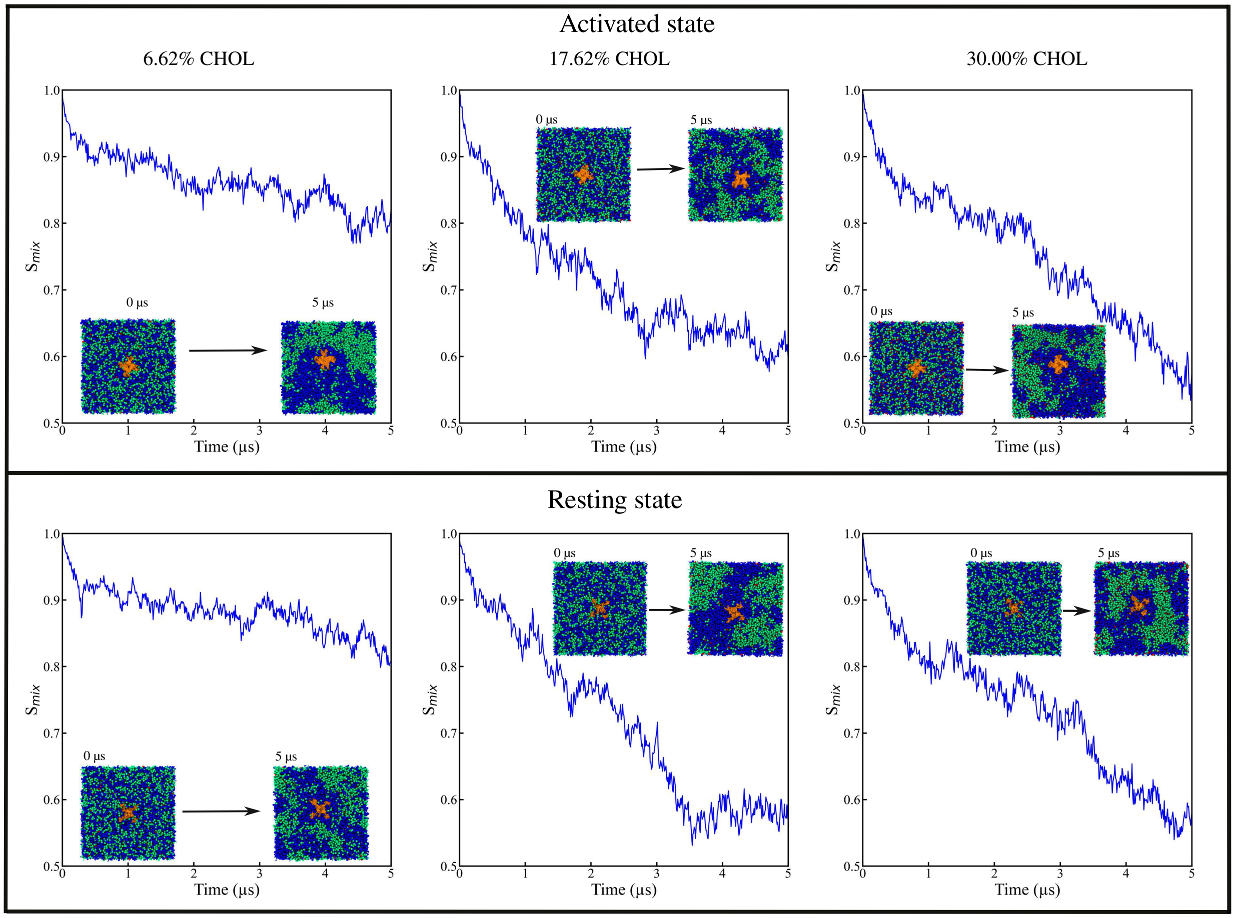


Figure S5: Time evolution of $S_{mix}$ for DIPC, DPPC and CHOL in activated (top row) and resting state NavABs (bottom row) for single channel NavABs at low, medium and high CHOL concentrations. Each plot include system snapshots at initial (0 $\mu$s) and final simulation time (5 $\mu$s) for NavAB systems. Color coding is same as in the main text.
